# Anti-SARS-CoV-2 swine glyco-humanized polyclonal antibody XAV-19 retains neutralizing activity against SARS-CoV-2 B.1.1.529 (Omicron)

**DOI:** 10.1101/2022.01.26.477856

**Authors:** Bernard Vanhove, Stéphane Marot, Gwénaëlle Evanno, Isabelle Malet, Gaëtane Rouvray, Françoise Shneiker, Edwige Mevel, Carine Ciron, Juliette Rousse, Pierre-Joseph Royer, Elsa Lheriteau, François Raffi, Odile Duvaux, Anne-Geneviève Marcelin, Vincent Calvez

## Abstract

B.1.1.529 is the SARS-CoV-2 variant designated Omicron by the WHO in November 2021. It is a highly divergent variant with a high number of mutations, including 26-32 mutations in the spike protein among which 15 in the Receptor Binding Domain (RBD) including at the human angiotensin converting enzyme 2 (ACE-2) receptor interacting interface. Because of a decreased affinity for the ACE-2 receptor and a geometric reorganization of the S1-S2 cleavage site, the Omicron variant is predicted to not have a significant infectivity advantage over the delta variant and to be less pathogenic than Delta. However, in Omicron, neutralizing epitopes are greatly affected, suggesting that current vaccines and neutralizing monoclonal antibodies might confer reduced protection against this variant. In contrast, we and others previously demonstrated that polyclonal antibodies against SARS-CoV-2 RBD obtained from hyperimmunized animal hosts do maintain their neutralizing properties against Alpha to Delta. Here, we confirmed these findings by showing that XAV-19, a swine glyco-humanized polyclonal antibody retains full neutralizing activity against Omicron.

## Introduction

Numerous neutralizing monoclonal antibodies have been developed against SARS-CoV-2. Most of them have been raised against the original Wuhan-type virus and see their neutralization potential abrogated or reduced against variants ^1^, especially against the Omicron variant. To anticipate these difficulties, polyclonal antibodies (pAbs) have been produced owing to their potential to bind multiple target epitopes and to maintain their neutralizing activity even though mutations occur in the antibody binding site ^2,3^. pAbs against SARS-CoV-2 of different sources are being used in at least 10 clinical trials (Table 1) but clinical proof of efficacy is still partial and limited to equine globulins ^4^. XAV-19 is a swine glyco-humanized polyclonal neutralizing antibody containing multiple immunoglobulins directed against a set of well-identified peptides of the Wuhan-type SARS-Cov-2 RBD protein ^5^. Glyco-humanization refers to the elimination of the animal-type α1,3-Gal and Neu5GC-Gal-β-1,4 GlcNac glycosylation epitopes by the human/ape counterparts Neu5AC-Gal-β-1,4-GlcNac epitopes. This is predicted to prevent allergies and serum sickness otherwise induced in 20 to 30% of the patients receiving animal-sourced immunoglobulins ^6,7^. Glyco-humanized polyclonal antibodies are produced from pigs where genes for the glycosylation enzymes α1,3-galactosyltransferase (GGTA1) and cytidine monophosphate N-acetyl hydroxylase (CMAH) have been inactivated ^8^. These animals are like those recently used in xenotransplantation trials ^9,10^. Recent clinical safety data confirmed that glyco-humanized swine pAbs are well tolerated and present better pharmacokinetic properties than other pAbs ^11^.

**Table 1:** pre-clinical and clinical-stage polyclonal antibody programs against SARS-CoV-2

| **Sponsor (country)** | **Type of polyclonal antibody (name)** | **Clinical trial reference and phase** |
| --- | --- | --- |
| Inmunova (Argentina) | Equine hyperimmune F(ab')<sub>2</sub> serum (INM005) | NCT04494984, phase 3 |
| Caja Costarricense de Seguro Social (Costa Rica) | Equine hyperimmune F(ab')<sub>2</sub> serum | NCT04838821, phase 2/3 |
| Hospital San Jose Tec de Monterrey/ Inosan Biopharma (Mexico) | Equine immunoglobulin F(ab')<sub>2</sub> fragments (INOSARS) | NCT04514302, phase 1 |
| Bharat serums and Vaccines (India) | Equine COVID-19 A F(ab')<sub>2</sub> antiserum (BSVEQAb) | NCT04834908, phase 2 |
| Butantan Institute (Brazil) | Equine COVID-19 F(ab')<sub>2</sub> antiserum | NCT04834089, phase 2 |
| Fab'entech (France) | Equine hyperimmune serum | Preclinical |
| National Institute of Allergy and Infectious Diseases/SAB Biotherapeutics (USA) | Humanized cattle globulins (SAB-185) | NCT04518410, Phase 3 |
| Nantes University Hospital/Xenothera (France) | Glyco-humanized pig globulins (XAV-19) | NCT04453384, Phase 2<br>EUDRACT 2020-005979-12, Phase 3 |
| GigaGen (USA) | Recombinant Hyperimmune Polyclonal Antibody (GIGA2050) | NCT04883138, Phase 1 |

We have previously described how XAV-19 binds multiple target epitopes on SARS-CoV-2 Spike and maintains neutralizing activity against the Alpha (United Kingdom/B.1.1.7), Beta (South African/B.1.351), Gamma (Brazil/P.1) and Delta (Indian/B.1.617.2) variants of concern. Here, we extended these observations to the variant Omicron (B.1.1.529). Using recombinant RBD protein of the Omicron type, we measured the neutralizing activity of XAV-19 against the Omicron ACE-2/RBD interaction in ELISA format and concluded that it is 100% conserved. In parallel the neutralization of the infectivity was confirmed in neutralization assays with live Omicron clinical isolate virus.

## Methods

### Reagents

XAV-19 is a swine glyco-humanized polyclonal antibody against SARS-CoV-2 produced by Xenothera by immunization of double knock out pigs for alpha 1,3-galactosyltransferase (GGTA1) and cytidine monophosphate N-acetyl hydroxylase (CMAH) genes, as previously described ^8^. Batch-to-batch consistency is ensured by the clonal nature of the immunized animals, thereby reducing the genetic diversity, by controlling the immunization process and applying specifications on the resulting serum, before it undergoes a downstream purification process. XAV-19 used in this study was the clinical batch DP1/B07-08. Comparator Evusheld^®^ (tixagevimab-AZD8895/cilgavimab-AZD1060) is from AstraZeneca (Cambridge, UK). Recombinant Spike molecules of the Wuhan type (Sino Biological ref 40591-V08H), mutation-containing RBD (Y453F ref 40592-V08H80; N501Y, ref 40592-V08H82; N439K, ref 40592-V08H14; E484K, ref 40592-V08H84), Alpha (ref 40591-V08H12; containing mutations HV69-70 deletion, Y144 deletion, N501Y, A570D, D614G, P681H) and Beta (ref 40591-V08H10; containing mutations K417N, E484K, N501Y, D614G) forms and recombinant human Fc-tagged ACE-2 were purchased by Sino Biological Europe, Eschborn, Germany. The mutation containing RBD corresponding to the Omicron variant (containing mutations G339D, S470L, S472P, S474F, K417N, N440K, G446S, S477N, T478K, E484A, Q493R, G496S, Q498R, N501Y, Y505H) purchased independently by Sino Biological Europe and by Covalab, France.

SARS-CoV-2 Wuhan (D614 and D614G B.1variant), Alpha, Beta, Gamma, Delta and Omicron strains were isolated from SARS-CoV-2 infected patients in the Pitié-Salpêtrière hospital, Paris, France.

### Binding ELISA

The binding of XAV-19 to Wuhan and Omicron RBD proteins was measured by ELISA as previously described (Vanhove et al, Front. Immunol. 2021). Briefly, recombinant RBD antigen (Sino Biological Europe, Germany) was immobilized on microtiter plate in carbonate/bicarbonate buffer pH 9.0 at 1μg/ml at 4°C for 16h. After washing and saturation with PBS-BSA-Tween, successive twofold dilutions of XAV-19 were added for 2h and washed. Porcine immunoglobulins were then revealed with a 1:1000 dilution of peroxidase-conjugated secondary antibody (Bethyl, USA). Binding intensity was revealed by addition of TMB reagent (Sigma, France) for 20 minutes, after which the reaction was stopped with 50 μl of 1M H2S04. The optical density was taken at 450 nm.

### Spike/ACE-2 neutralization assay

Neutralization of the interaction of SARS-CoV-2 RBD with its receptor ACE-2 was also measured by ELISA as previously described ^5^. Briefly, recombinant SARS-CoV-2 Spike S1-HIS proteins (Sino Biological Europe for the Wild Type Wuhan, or Omicron from Sino Biological Europe and from Covalab, France, for an alternative source of the Omicron form) were immobilized on microtiter plate in carbonate/bicarbonate buffer pH 9.0 at 1μg/ml at 4°C for 16h. After washing and saturation with PBS-BSA-Tween, successive dilutions of XAV-19 were added for 30 min, followed by ACE-2-mFc Tag ligand (Sino Biological; final concentration 125 ng/ml). After incubation for 1h at room temperature and washing, the ACE-2 mFc Tag was revealed using a peroxidase-conjugated anti-mouse secondary antibody (1:1000 dilution). Binding intensity was revealed by addition of TMB reagent (Sigma, France) for 6 min, after which the reaction was stopped with 50 μl of 1M H2S04. The optical density was taken at 450 nm.

### Cytopathogenic Effect (CPE) assay

Vero cells (CCL-81) and Vero E6 cells (CRL-1586) were obtained from the American Type Culture Collection and maintained at 37°C with 5% CO2 in Dulbecco’s Modified Eagle’s Medium (DMEM), supplemented with 5% heat-inactivated fetal bovine serum (FBS) and 1X Penicillin-Streptomycin solution (Thermo Fisher Scientific, USA). SARS-CoV-2 clinical isolates (D614G variant; GenBank accession number MW322968), Alpha (GenBank accession number MW633280), Beta (GenBank accession number MW580244), Gamma (Gene accession number pending), Delta (Gene accession number pending) and Omicron (Gene accession number pending) were isolated from SARS-CoV-2 RT-PCR confirmed patients by inoculating Vero cells with sputum sample or nasopharyngeal swabs in the biosafety level-3 (BSL-3) facility of the Pitié-Salpêtrière University Hospital. Viral stocks were generated using one passage of isolates on Vero cells. Titration of viral stock was performed on Vero E6 by the limiting dilution assay allowing calculation of tissue culture infective dose 50% (TCID50). The neutralizing activity of XAV-19 was assessed with a whole virus replication assay using the five SARS-CoV-2 isolates. XAV-19 was subjected to serial two-fold dilution ranging from 50 μg/ml to 0.05 μg/ml in fresh medium. 50 μl of these dilutions were incubated with 50 μl of diluted virus (2 × 10^3^ TCID_50_/ml) per well in a 96-well plate at 37°C for 60 min in 8 replicates. Hundred μl of a Vero E6 cell suspension (3 × 10^5^ cells/ml) were then added to the mixture and incubated at 37°C under an atmosphere containing 5% CO_2_ until microscopy examination on day 4 to assess cytopathogenic effect (CPE), as previously described (Vanhove et al, Frontiers Immunol. 2021). For viral load (VL) quantification, a similar experiment was conducted with a range of XAV-19 dilution ranging from 24 μg/ml to 1 μg/ml in fresh medium. On day 4, RNA extraction of the 8 pooled replicates of each XAV-19 dilution was performed with NucliSENS EasyMag (BioMerieux) according to manufacturer’s protocol. The relative VLs were assessed from cycle threshold values for ORF1ab gene obtained by the TaqPath™ COVID-19 RT-PCR (ThermoFisher, Waltham, USA) and by linear regression in log10 copies/ml with a standard curve realized from a SARS-CoV-2 positive nasopharyngeal sample quantified by Droplet-Digital PCR (Bio-Rad). IC50s were analyzed by nonlinear regression using a four-parameter dosage-response variable slope model with the GraphPad Prism 8.0.2 software (GraphPad Software, USA).

## Results

### XAV-19 has target epitopes lying outside the Omicron mutations sites

We previously determined by peptide mapping and proteolytic epitope mapping that antibodies contained in XAV-19 recognize virtually all peptides present in the Wuhan SARS-CoV-2 RBD but that the dominant target epitopes lied in 4 regions underlined in Figure 1 (347-fasvyawnr-417, 409-qiapgqtgn-417, 445-vsgnynylyrlfrksnlkpferdisteiy-473, 530-stnlvk-535). Interestingly these regions contained 6 amino acids on the 17 described to be directly involved in the contact with ACE-2 counter receptor ^12^(outlined in blue in Figure 1), probably explaining the neutralizing potency of XAV-19. Among the 15 Omicron mutations contained in RBD (highlighted in red in Figure 1), 7 amino acids belong to those important for the contact with ACE-2 and only 2 of them lie inside the major XAV-19 target epitopes (N417 and S446). In comparison, 5 amino acids belonging to the target epitope of tixagevimab and cilgavimab correspond to amino acids mutated in Omicron (dotted underlined; K440, S446, N447, K478, A484, according to Dong et al ^13^.

**Figure 1.**
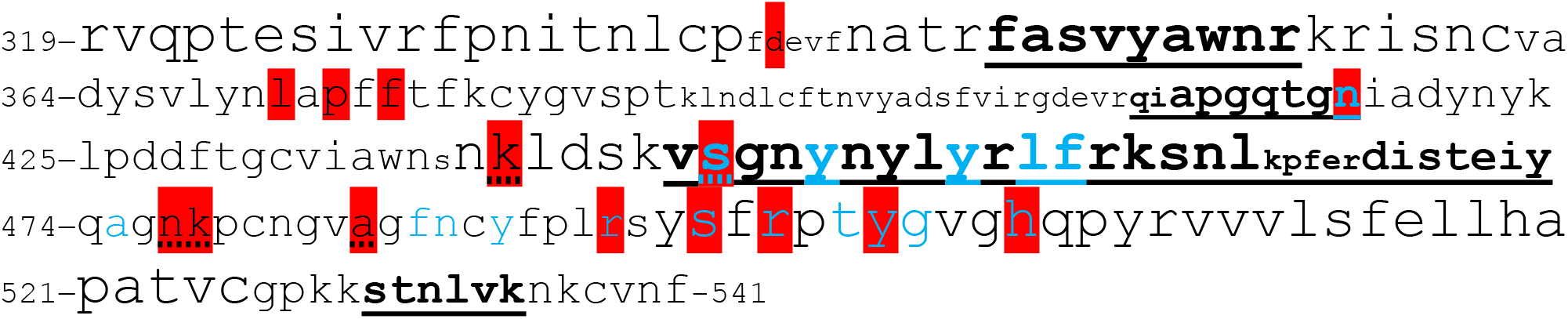
Amino acid sequence of the SARS-CoV-2 spike RBD variant Omicron and XAV-19 target epitopes (amino acid sequence numbered according to DBSOURCE sequence reference NC_045512.2). As previously described, all peptides are potential targets of XAV-19 antibodies as measured by peptide scanning, with binding intensity reflected by the font size (small font, weak binding to the corresponding peptides; medium font, medium binding; large font, strong binding). Underlined epitopes are XAV-19 target epitopes also confirmed by proteolytic epitope mapping (Vanhove et al, Front. Immunol.). Blue amino acids: amino acids in contact with ACE-2, according to Jafary et al. ^28^. Amino acids highlighted in red: mutations found in Omicron, differentiating from the original Wuhan RBD.

### XAV-19 binding to SARS-CoV-2 Omicron Spike

The binding intensities of XAV-19 against RBD Wuhan and Omicron were compared by ELISA. Although some differences appear in the profiles, the results indicated that XAV-19 binding reached the same plateau at similar concentrations (approximately 1 μg/ml).

### Neutralization of the interaction of Omicron Spike with ACE-2

XAV-19 was tested in a RBD/ACE-2 binding competition assay, where the RBD protein was of the original Wuhan type or contained the mutations corresponding to the Omicron variant. Two independent sources of recombinant Omicron RBD were used. The dose response profiles were similar whether Wuhan or Omicron RBD were used, indicating similar full neutralization potency of XAV-19 against their interaction with ACE-2 (Figure 3A). In the same assay, the combination of tixagevimab and cilgavimab (Evusheld^®^) was assessed in parallel and revealed an efficacy limited to approximately 60% neutralization, whatever the dose (Figure 3B).

**Figure 2.**
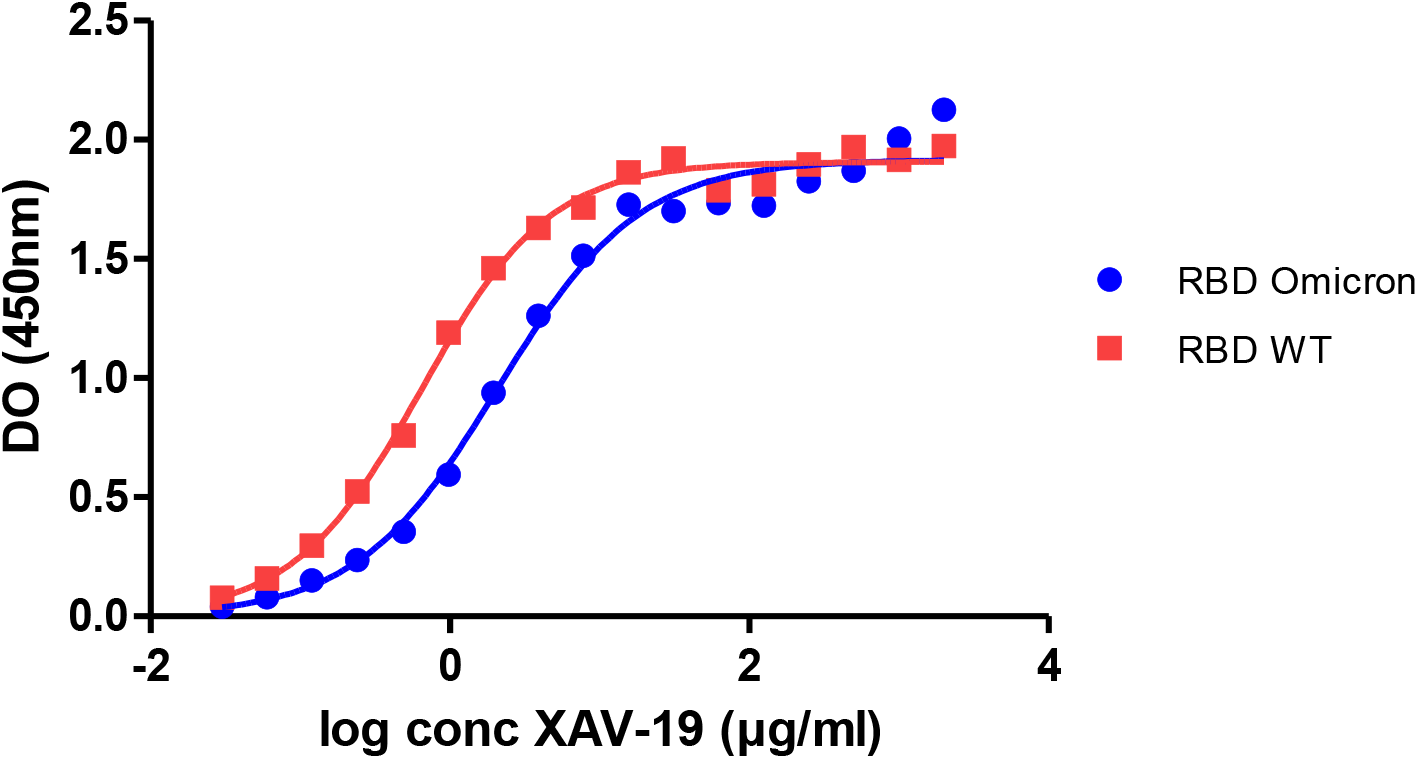
Binding intensity of XAV-19 to RBD Wuhan versus Omicron. Microtiter plates have been similarly coated with Wuhan or RBD recombinant proteins. XAV-19 was added at the indicated concentration and after washing revealed with a peroxidase-conjugated secondary antibody. Optical density (means of triplicate measurements run in a single experiment) was recorded after addition of TMB reagent.

**Figure 3.**
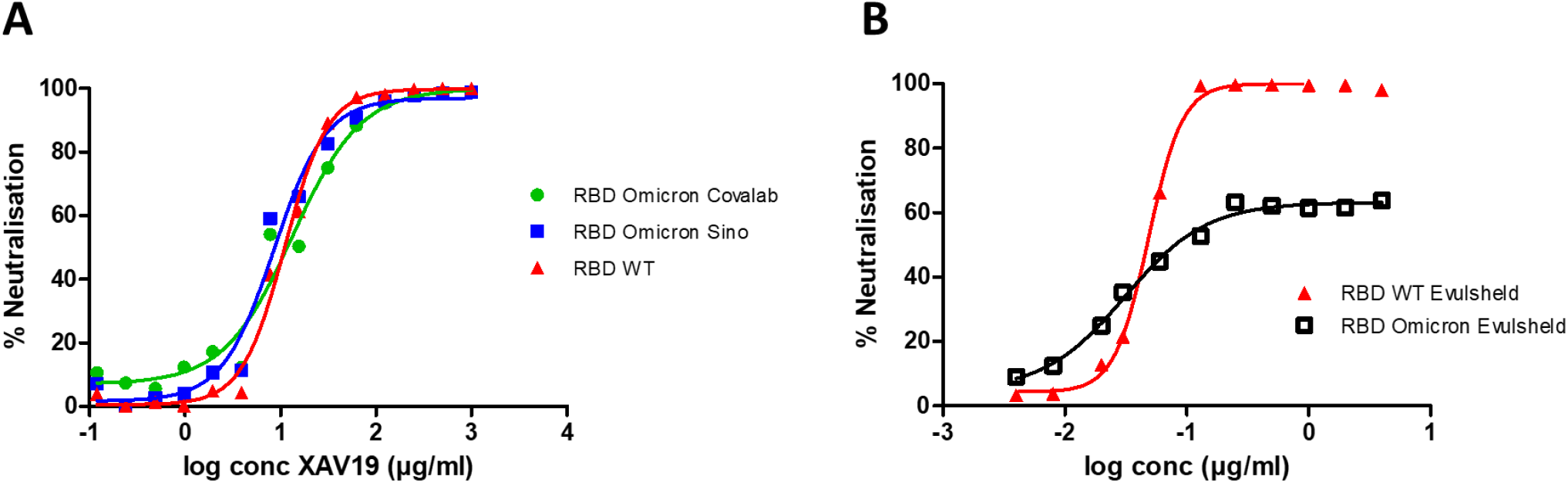
Neutralization assay in the ELISA format: assessment of SARS-CoV-2 Spike/ACE-2 interaction and its XAV-19-mediated inhibition. Spike-HIS corresponding to the Wuhan or the Omicron (two different sources, as indicated) variants was immobilized on plastic and binding of recombinant human ACE2-Fc was revealed with a secondary antibody against Fc. 100% inhibition represents absence of Spike/ACE-2 interaction. A: Data are means of triplicate measurements run in a single experiment assessing XAV-19 at the indicated concentration. B: Means of triplicate measurements run in a single experiment assessing a mixture of tixagevimab and cilgavimab at the indicated global concentration.

**Figure 4.**
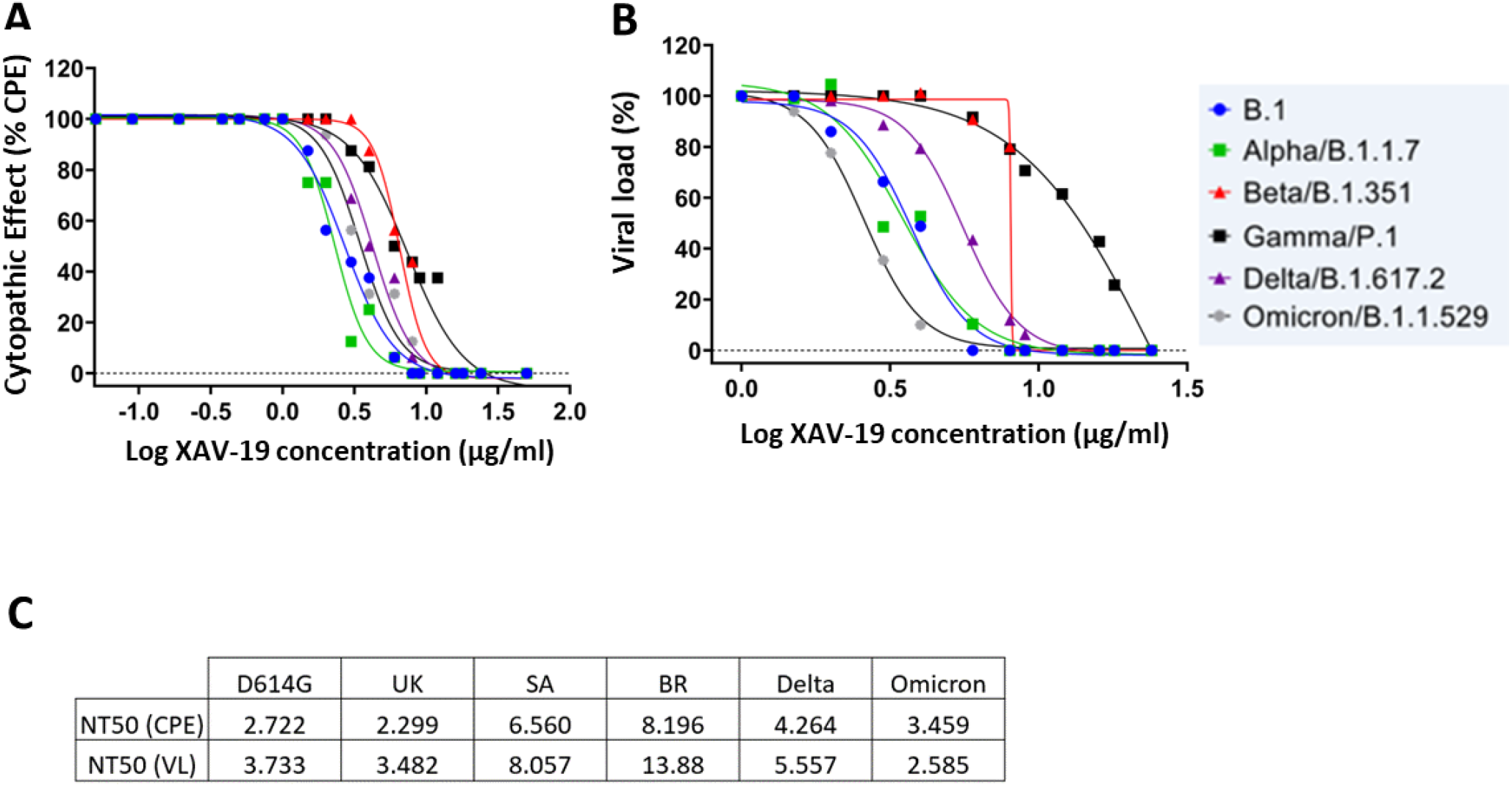
In vitro neutralization of SARS-CoV-2 variants by XAV-19. XAV-19 neutralizing potency was evaluated in an in vitro assay using whole replicating viruses. Percentage of infection was evaluated as described in Materials and Methods, based on cytopathogenic effect (CPE) (A) and virus RNA load (B) after infection with SARS-CoV-2 viruses of the indicated variants. CPE percentage was assessed by microscopy examination and calculated on 8 replicates for each XAV-19 concentration. 100% represent absence of CPE inhibition at the studied concentration, as found in control (no inhibitor) condition. Viral load percentage was calculated as the ratio of viral load in each XAV-19 concentration to viral load in controls (no inhibitor). XAV-19 concentrations are expressed on a 10 logarithmic scale.

### Neutralization of live SARS-CoV-2 Omicron variant

The neutralization of viral infectivity of XAV-19 was determined by CPE assays using Wuhan, Alpha, beta, Gamma, Delta and Omicron SARS-CoV-2 clinical isolates, as previously described ^14^. The test assessed inhibition of live viruses with sensitive Vero E6 cells and recorded infection after 4 days by measuring CPE and viral load by RT-qPCR. As previously shown ^5^, XAV-19 neutralized 100% of CPE or reached 100% viral load reduction with slightly varying NC50 concentrations. Interestingly, NT50 against Omicron was in the low range in CPE and lower than for all other variants in viral load assessment.

## Discussion

Data published in 2021 ^5^ indicated that for XAV-19, an ELISA binding test of RBD to the human ACE-2 receptor was predictive of the neutralizing activity measured in a neutralization assay with live virus and sensitive cells. We have now measured the neutralizing activity of XAV-19 against the RBD Omicron variant by ELISA and concluded that a full neutralization was reached at a dose similar to that obtained with the RBD of the original Wuhan virus. As a comparator, we used a combination of tixagevimab and cilgavimab (Evusheld^®^) in the same ELISA and observed a limited neutralization, which might be attributed to amino acid changes in the target epitopes of these monoclonal antibodies. Changes between Wuhan and Omicron RBD in the target epitopes of cilgavimab represent 16% of the amino acids (3 changes on 19 amino acids) and 21% for tixagevimab (3 changes on 14 amino acids). By overlying XAV-19 target epitopes, Omicron mutations and amino acids described important for the contact with ACE-2 (as in Figure 1), it was clear that most Omicron mutations lied mostly outside the target epitopes: changes between Wuhan and Omicron RBD represents only 3% of the main target epitopes of XAV-19 (2 changes on 53 amino acids). This might justify that binding of XAV-19 to Omicron RBD was mostly preserved. This might also justify the full neutralizing activity of XAV-19 against live SARS-CoV-2 Omicron in infection assays at equivalent to lower concentration than against other variants.

Fantini et al ^15^ reported that Omicron presents a decreased affinity for the ACE-2 receptor, probably as a result of the alteration of several amino acids important for direct contact with ACE-2. If XAV-19 binds similarly to RBD Omicron which itself has a weaker binding to ACE-2, the model predicts that XAV-19 should present a preserved or higher neutralization potency against Omicron than against any other variant. This prediction was verified experimentally by ELISA and CPE assays on sensitive Vero E6 cells, where the IC50 were similar or even better when viral load was measured.

Most monoclonal antibodies developed against SARS-CoV-2 RBD of the Wuhan form see their neutralizing efficacy abrogated or strongly impacted against the Omicron variant ^16^. This is probably due to the high mutation burden in the RBD that has a high probability of impacting monoclonal antibodies target epitopes. Having multiple alternative target epitopes on the same target, as previously shown ^5^, makes it less likely for a pAb to lose binding capacity even with such a high mutation burden. That most important target epitopes of XAV-19 lied in domains where Omicron mutations are absent is also a feature explaining the current observation that binding and neutralization properties are preserved.

COVID-19 clinical trials with at least 10 different pAb preparation are ongoing, among which XAV-19 (phase 3 clinical trial NCT0492830/EUDRACT 2020-005979-12). As for mAbs, pAbs should be administered to COVID-19 patients early enough in their disease course so that the viral neutralization impacts the clinical status. Provided patients responsive to antibody treatment can be identified in such trials, pAbs and XAV-19, because they broadly neutralize variants, might represent a robust therapeutic alternative to mAbs to fight against Omicron and possibly other future variants.

## Key words

Polyclonal antibodies, pig, Covid-19, SARS-CoV-2, Spike, neutralization, variant of concern, Omicron

## Authorship

Conceived the study: SM, BV, VC, IM, AGM, OD, FR

Designed and supervised some experiments: SM, BV, IM, VC, AGM, EM, OD, BV, GE, EL, GR

Performed the experiments: SM, GE, CC, JR, PJR

Analyzed data and wrote the manuscript: BV, OD, VM, FS

## Acknowledgments

This work was supported by Xenothera, the Agence Nationale de la Recherche sur le SIDA et les Maladies Infectieuses Emergentes (ANRS MIE), AC43 Medical Virology and Emergen Program, the SARS-CoV-2 Program of the Faculty of Medicine of Sorbonne Université and by Bpifrance, grant « Projet de Recherche et Développement Structurant Pour la Compétitivité spécifique à la crise sanitaire COVD-19 – POLYCOR».

